# Floral herbivory does not reduce pollination-mediated fitness in shelter rewarding Royal Irises

**DOI:** 10.1101/184382

**Authors:** Mahua Ghara, Christina Ewerhardy, Gil Yardeni, Mor Matzliach, Yuval Sapir

## Abstract

Florivory, the damage to flowers by herbivores can affect fitness both directly and indirectly. Flowers consumed by florivores may fail to produce fruit or produce lower seed set because of direct damage to reproductive organs. In addition, eaten flowers are less attractive to pollinators because of reduced or modified advertisement, which reduces pollination services. While observational data are abundant, experimental evidence is scarce and results are contrasting. We tested experimentally the effect of florivory on both pollinator visitation and reproductive success in three species of the Royal Irises, which have large flowers that are attractive to pollinators, and potentially also for florivores. We hypothesized that florivory will reduce pollen deposition due to reduced attractiveness to pollinators, while fruit set and seed set will depend on the extent of florivory. We performed artificial florivory in two experiments over two years. In the first experiment, each of the three floral units of a single *Iris* flower was subject to either low or high artificial florivory, or left un-touched as control. We counted the number of pollen grains deposited on each of the three stigmas as a measure of pollinator visitation. In the second experiment, three flowers of the same plant received low, high, or no artificial florivory and were further recorded for fruit and seed production. In 2016, high artificial florivory revealed lower number of pollen grains on stigmas of *Iris atropurpurea*, but in 2017 there was no difference. Similarly, number of pollen grains in high artificial was lower than low florivory in 2017 in *I. petrana*. No significant effect of florivory was found on pollen grain deposition, fruit set or seed set. The results remained consistent across species and across years. The results undermine the assumption that flower herbivory is necessarily antagonistic interaction and suggests that florivores may not be strong selection agents on floral reproductive biology in the *Oncocyclus* irises.

## INTRODUCTION

Flowers of animal-pollinated plants are the major means of plants to advertise and attract pollinators. Floral traits serve as signals to the pollinators, usually fit to the most-efficient pollinator (Fenster et al., 2004). Flowers advertise through visual as well as olfactory signals, and even by acoustic signature, in order to stand out of the canopy or other flowering species (Schiestl and Johnson, 2013). Consequently, flower traits that increase attraction are selected by pollinators (Harder and Johnson, 2009). For example, floral size, contributing to the visibility of the flower, is positively selected by pollinators (Sletvold et al., 2010, 2016; Campbell et al., 1991; Conner and Rush, 1996; Harder and Johnson, 2009; Lavi and Sapir, 2015). However, flowers are costly organs that require investment in resources for production and maintenance. Other selection agents also act on floral traits, either in concert with, or in contrast to the selection exerted by pollinators (Strauss and Whittall, 2006). For example, floral size is under contrasting selection regimes, positive by pollinators and negative by drought and water loss (Galen, 2000; Carroll et al., 2001). Color polymorphism is also thought to be maintained by the combined effect of mutualists (pollinators) and antagonists (Carlson and Holsinger, 2010), or by opposing selection directions by herbivores and pathogens (Frey, 2004). Although many studies have examined floral adaptations to pollinators, the role of non-pollinator selection agents in shaping floral evolution and plant reproductive success is still understudied.

Florivory, the damage of flowers by herbivores, is widespread across plant taxa and ecosystems (Gonzáles et al., 2016; Burgess, 1991). Florivores comprise of various taxonomic orders of animals that consume the entire flower (or the buds) or floral parts (bracts, sepals, petals, nectaries, stamens, pistils or pollen). Florivory may affect plant fitness directly or indirectly, by reducing fruiting or seed-set, and can consequently affect population dynamics (Louda and Potvin, 1995). Florivory reduces fitness directly when reproductive parts are consumed, whereas indirect effect can be through effects on pollinator behavior and hence reduced pollination services. An increasing number of studies suggest that florivory can decrease pollinator visitation rate and pollination success (reviewed in Gonzáles et al., 2016, but see Zhu et al., 2017).

Methods for studying the effect of florivory and its consequences on pollination primarily focus on the effect of florivore presence or the extent of florivory on pollinator visits, using natural encounters and manipulative studies. For example, Kirk et al. (1995) placed black spots on flowers to simulate presence of florivore beetle and found that bees were likely to avoid flowers with mimicry of florivore presence. Although artificial florivory performed in the field can provide an estimate to the direct effect of florivory on pollination, few studies implement this method. Moreover, behavior of pollinators and/or measure of maternal fitness (measured as the number of visits or seed set, respectively) are often used to estimate the effect of florivory on pollination (e.g., Cardel and Koptur, 2010).Pollen deposition on stigma can be a direct evidence for pollination success, but has rarely been incorporated in studies that investigate the effect of florivory on pollination. Therefore, controlled manipulative experiments that measure pollen deposition and fitness are needed to estimate directly the interaction of florivory and pollination success.

In a survey of the literature, we found only a small number of studies that used controlled experimental florivory that mimics natural florivory by artificially manipulating or removing parts of the petals and tested for effect on pollination. Of these, seven studies found negative effect on pollination success, one study found mixed effects that depended on flower morph (Carper et al., 2016), and one study found no effect (Tsuji et al., 2016). Thus, it remains unclear whether florivory affects plant fitness indirectly via negative effects on pollinators. As an example, McCall (2008) found that both natural and artificial petal damage indeed reduced fitness, and while it deterred pollinator activity, the effect was a result of petal physical damage rather than reduced pollinators’ activity, as pollen addition treatment did not recover fruit-set. Hence, for fully understanding the multiple effects of florivory on fitness and the interplay between direct and indirect effects it requires further experiments in diverse plant species.

We studied the effect of florivory on pollination success in three species of the Royal Irises *(Iris* section *Oncocyclus*) in their natural habitats in Israel (Fig. 1 A-C). The Royal Irises comprise about 30 species across the Middle East, and most are narrow endemics (Rix, 1997; Mathew, 1989). This group of irises is well studied as a system for pollination by shelter reward to male *Eucera* bees (Fig. 1 D; Sapir et al., 2005; Sapir et al., 2006; Watts et al., 2013; Monty et al., 2006; Vereecken et al., 2013; Lavi and Sapir, 2015). Flowers of the Royal Irises grow singly on a stem, but a plant (genet) comprises one to hundreds of stems (ramets) in a well-defined patch. Flowers of the Royal Irises comprise of three identical units, each bearing one upright and one pendant petal (standard and fall, respectively), and one petaloid style, curved above the fall to create a tunnel where the reproductive organs reside (Fig. 1). The three stigmas, located at the top of the entrance of each of the pollination tunnels. Pollen is deposited on stigmas by bees that move among flowers as they seek shelter (Sapir et al., 2005). The three style lobes are merged at the base to form one united style. A previous study showed that pollinating one style is sufficient to produce seeds in all three carpels in the ovary (Watts et al., 2013), thus, although each pollination tunnel functions as a single ecological unit of pollination, pollinator visit in one tunnel affects the fitness of the whole flower. All the species are self-incompatible and require pollination to set fruits (Sapir et al., 2005).

**Figure 1.**
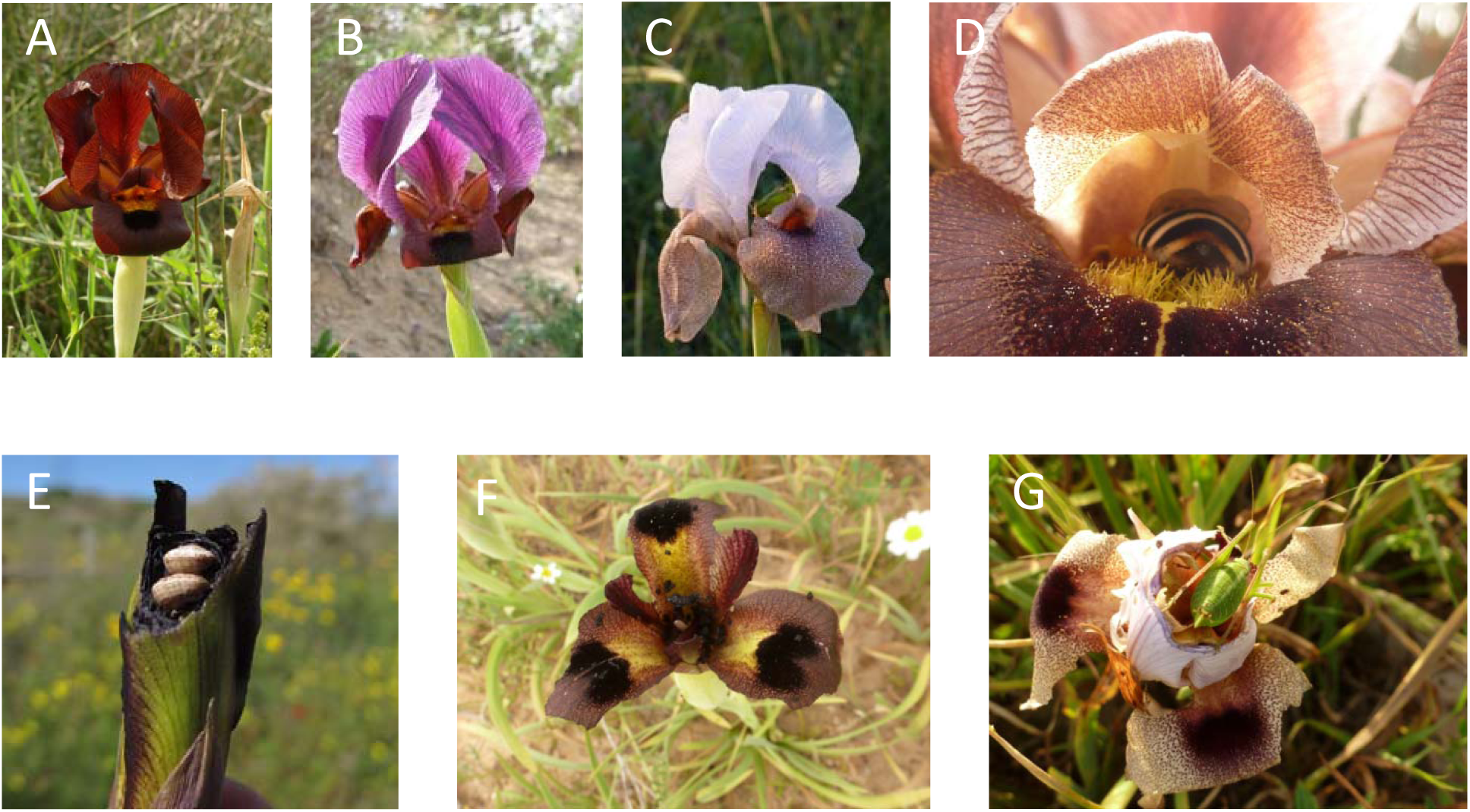
(A) *Iris atropurpurea* in Netanya; (B) *Iris petrana* in Yeruham; (C) *Iris lortetii* in Malkiya; (D) Male *Eucera* bee, the specific pollinator of the Royal Irises, sheltering within a pollination tunnel of *Iris petrana*; (E-G) Natural florivory in flowers of *Iris atropurpurea* (E), *I. petrana* (F) and *I. lortetii* (G).

During a survey in the year 2015 and also based on previous observations we found that the Royal Irises are eaten by various florivores, from snails and true bugs to grasshoppers, birds and goats, and the intensity of damage ranges from a few superficial scratches or poke marks, to >90% of damage to floral tissue (M. Ghara and Y. Sapir, unpubl. Res.; Fig. 1 E–G). From our observations, it appears that all flower parts are potentially eaten, including tepals, the petaloid style, anthers and ovaries. While damage to reproductive organs may obviously reduce fitness directly, we asked whether florivory of the advertising floral parts affects fitness indirectly through reduced attraction to pollinators. In order to estimate the effect of florivory on pollinator visitation and pollination success, we manipulated the flowers to simulate two levels of florivory, i.e., high (more than 50% damage) and low (10-30% damage), and compared them to control flowers without damage. We asked the following questions: (1) Does florivory affect pollinator visitation? (2) Does florivory affect fruit set and seed set? We used pollen deposition on the stigma as a surrogate for estimating pollinator visitation and used fruit- and seed-set to estimate overall effect of florivory on (maternal) fitness. Accounting for two measures of pollination success simultaneously is likely to reveal an indication of the possible effect of florivory on pollination-mediated fitness in the Royal Irises.

## MATERIALS AND METHODS

### Study species and sites

Three species of the Royal Irises (Iridaceae), *Iris atropurpurea* Baker, *I. petrana* Dinsm., and *I. lortetii* Barbey ex Boiss, were used in this study. Experiments were conducted in the natural environment at the largest population for each of the species. *Iris atropurpurea* was studied in Netanya Iris Reserve (32.28°N, 34.84°E, alt. 35 m), located on stable coastal sand dunes in Mediterranean climate and consisting of mostly low shrub vegetation. Population size is estimated >1,000 plants (Yardeni et al., 2016). Flowering season, and hence experiment time, is earlier compared to other species of the Royal Irises, starting as early as mid-January, and peaks in February. Experiments in *I. atropurpurea* in Netanya were conducted between 12 February 2016 and 02 March 2016, and between 19 February 2017 and 07 March 2017. *Iris petrana* was studied in Yeruham Iris Reserve (31.02°N, 34.97°E, alt. 560 m), a large population (estimated >10,000 plants) growing on sandy loess hills over Neogene sandstone in arid climate. Vegetation is sparse desert shrubs, mostly *Retama raetam* (Forssk.) Webb and *Anabasis articulata* (Forssk.) Moq. Flowering season is in March, and experiments in *I. petrana* were conducted between 05 March and 14 March in 2016 and between 19 March and 02 April in 2017. The shift in dates in 2017 resulted due to a delay in flowering period by about two weeks. *Iris lortetii* was studied in two sub-populations near Malkiya in the upper Galilee (central coordinates: 33.09°N, 35.52°E, alt. 620 m). Populations of *I. lortetii* are sparse and relatively small, thus, two sub-populations at a distance of 3 km of each other were pooled to achieve a sufficient sample size. Plants are growing on Eocene limestone in mesic Mediterranean climate and vegetation is open woodland dominated by *Quercus calliprinos* Webb and *Pistacia atlantica* Desf. trees, accompanied by dense herbaceous vegetation. *Iris lortetii* is the late blooming species among the Israeli species of the Royal Irises; experiments were conducted between 29 March and 07 April in 2016. The experiments were not conducted on *Iris lortetii* in 2017 because of high rhizome herbivory in 2016 and therefore a potential decrease in sample size.

Plants for the experiments were randomly selected in a dense part of the population in Netanya, or along transects in Yeruham. In Malkiya the plants are sparse and plants in all genets located were used. The three experiments described below were conducted simultaneously in time with only a single experiment conducted in each genet to avoid the joint effect of several treatments.

### Florivory manipulations and effect on pollination

To study the effect of florivory on pollen deposition, a single bud in a genet was bagged with a cloth bag and upon anthesis the bag was removed and each flower unit was given one of three treatments. In each treatment, the fall treated was paired with the standard in the opposite side of the flower. This way, a putative pollinator approaching the flower, assuming it is approaching in perpendicular angle to the pollination tunnel, is seeing the projection of the fall in the proximate part of the flower and the standard in the distant part. It is important to note that a confounding effect of such design is that the flower’s symmetry is changed. However, this design is advantageous because it enables to control for differences among flowers in, e.g., size of the pollination tunnel or colour. We employed two florivory treatments in each flower: High damage –the lower petal and its opposite upper petal were manually damaged up to 50% or more of their area, pierced by an awl of 6-8 mm in diameter. Low damage – 10-30% of the petals’ area were pierced using awl. The third pair of petals was left un-touched as control (Fig. 2).

**Figure 2.**
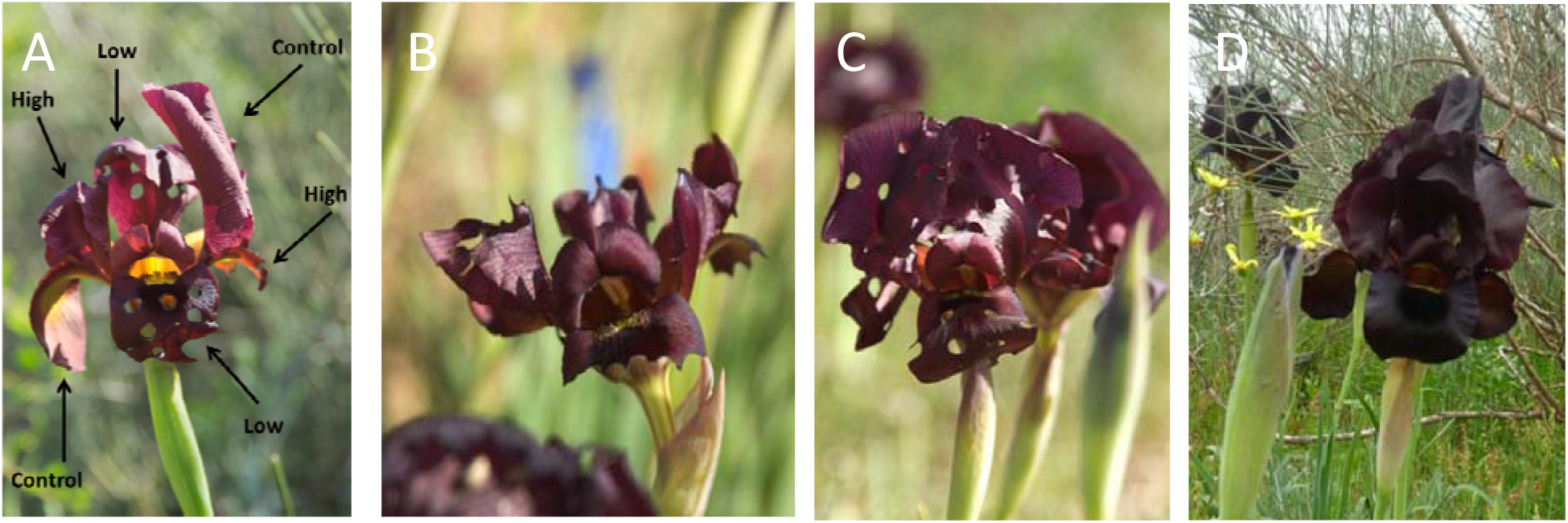
Artificial florivory manipulations exemplified in *Iris atropurpurea*. (A) Within flower manipulation – each floral unit treated as either high florivory (>50% petal cut), low florivory (10-30% petal removed by hole puncher), or control (no treatment). These treatments were used for testing the effect on pollination. (B–D) Flowers used for testing the effect of florivory on fitness. (B) Flower treated as high florivory. (C) Flower treated as low florivory. (D) Control flower.

To control for the possible effect of the contact between the metal awl and the flower the awl was rubbed on the petal surface in the control treatments. In addition, because damaging the petals required holding a layer of tissue paper against the awl, we also gently rubbed tissue paper under the surface of the petal in the control treatment. Note that florivory was done on the petals and not on the pollination tunnel, to explicitly address the hypothesis that change in visual display disrupt pollination services in the presence of florivory. Flowers that had natural florivory were not used. To prevent naturally occurring florivores from damaging the flowers, the stem of the flower was coated with a layer of double sided sticky paper tape, as well as a layer of Petroleum jelly (Vaseline). Occasionally we found insects trapped on the Vaseline layer, and in some rare cases, we found florivores that passed this barrier mainly because the sticky tape lost its stickiness owing to sand in the air that got stuck to the tape. Flowers found to be damaged naturally (mostly by flying insects, snails, or mammals) were discarded in order to account for the effect of controlled artificial florivory only.

After manipulating the flowers they left open for two consecutive evenings to enable pollinators to visit naturally. On the morning of the third day, the entire stigmas of the three pollination units were collected in vials containing 1 ml of 70% ethanol. Collected stigmas were brought to the laboratory for pollen counting and kept at room temperature. The vial containing the stigma with pollen was washed until there was no remaining pollen in alcohol. This was to ensure that pollen was not lost due to the protocol followed. Pollen grains were stained using a drop of basic fucshin (Calberla's stain). Each stigma was then dissected in a drop 70% aqueous glycerol (Dafni et al., 2005), mounted on microscope glass slide, and the pollen present on the stigma was counted under dissecting microscope (WILD Heerbrugg Switzerland M5-72558).

### Florivory manipulations and effect on maternal fitness

To study the effect of florivory on seed set, three buds of the same genet, roughly of the same developmental stage (i.e., before emerging from bracts) were selected and bagged to avoid bud florivory. Upon anthesis, each of the three flowers was randomly assigned to one of the florivory treatments described above, i.e., high, low, or control. To control for the effect of visual attraction mediated by flower size (Lavi and Sapir, 2015) we measured flower length as a surrogate for display size of each flower before manipulation. Flower length was measured from the bottom of the lower petal to the top end of the upper one. The flowers were left open to enable naturally occurring pollination. At the end of the season, approximately three weeks after the end of the flowering in each site, the fruits were collected and brought to the laboratory. In 2016, fruits of *I. atropurpurea, I. petrana* and *I. lortetii* were collected in 21 March, 02 April and 27 April, respectively. In 2017, fruits were collected in 12 March and 22 March for *I. atropurpurea* and on 13 April and 27 April for *I. petrana*. Fruits were kept in paper bags at room temperature until seed maturation. Fitness was recorded as presence or absence of a fruit (binomial data), and as the number of viable seeds (count data).

### Data analyses

Data were analyzed in R (R Development Core Team, 2014) using R-studio interface. Because each stigma is not independent of the other two stigmas of the same flower, we normalized the number of pollen grains on stigma by subtracting the number from the mean number of pollen grains on the three stigmas of the same flower. These normalized values of number of pollen grains deposited on stigmas were analyzed using generalized linear models (GLMs) with year and treatment effects nested within species (note that in *I. lortetii* experiment was done only in 2016). For analyses using fruit or seeds as explained variables, we used GLM and incorporated flower size as a covariate. For seeds as a response variable, we used only the subset of flowers that set a fruit. In order to account for non-normal distribution, we used GLM with binomial distribution errors for fruits and GLM with quasi-Poisson distribution errors for seeds.

## RESULTS

### Pollen deposition on stigma

Using absolute number of pollen grains deposited on the stigma, we found significant differences between species and years (GLM with quasi-Poisson distribution; *F_2,685_*=4.39, *P*=0.013 for species and F*_2,685_*=25.09, *P*<0.001 for year effect, nested within species). In *Iris atropurpurea*, 62 and 44 flowers were manipulated in 2016 and 2017, respectively. Of these, 49 (26.3%) of the stigmas did not receive pollen at all in 2016 and 8 (5.3%) in 2017. Flower units with artificial florivory, either high or low, received lower number of pollen grains, compared to control, un-manipulated units in 2016 (Contrast analysis: P=0.024), but the difference in number of pollen grains between high and low damage was not significant (*P*=0.115; Fig. 3 A). In 2017 higher number of pollen grains was deposited on stigmas overall, but with no differences among treatments (*P*=0.65; Fig. 3 B).

**Figure 3.**
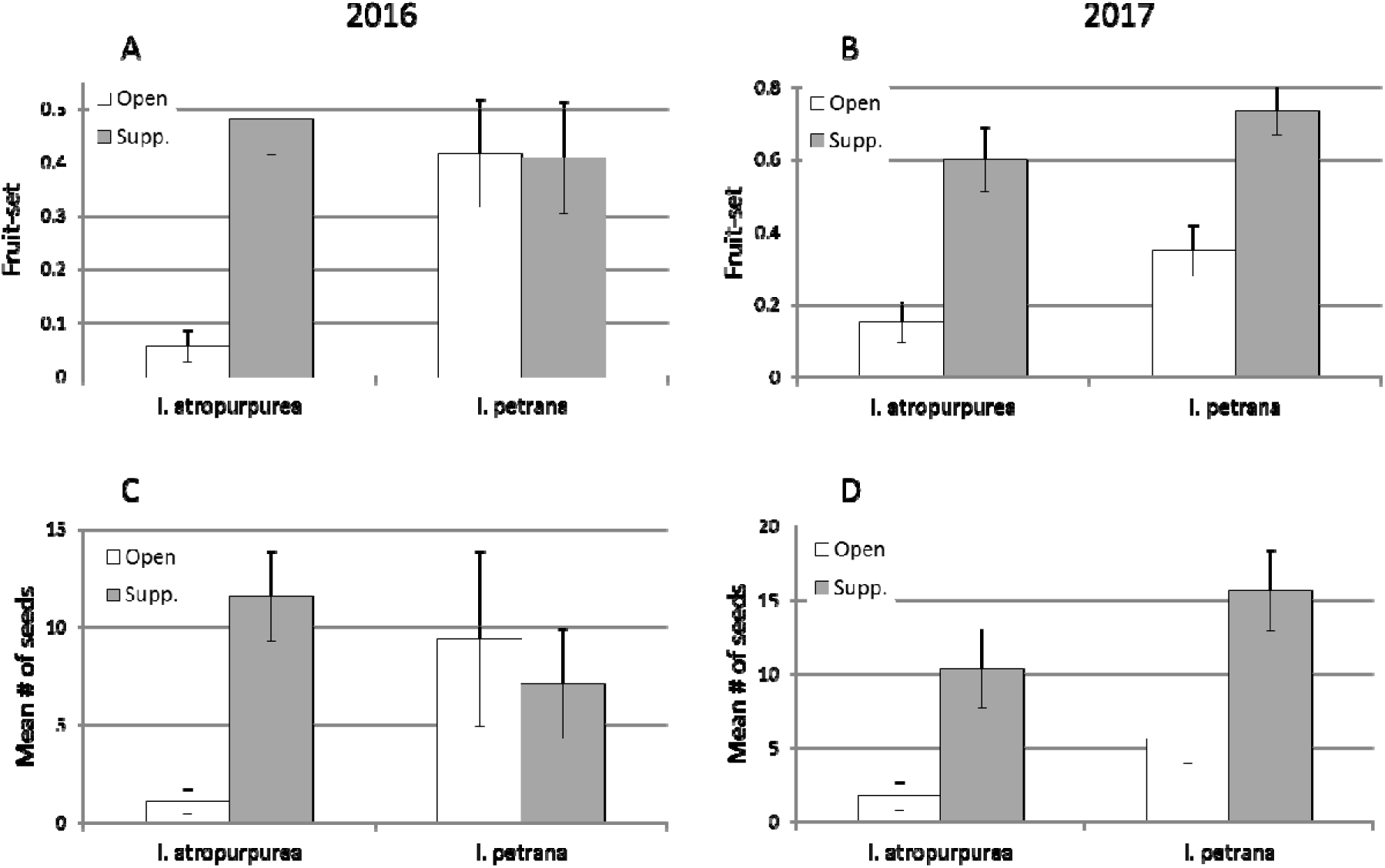
Mean number of pollen grains counted on stigma (± standard errors) as a function of florivory treatment in three species. (A) Pollen grains in 2016 experiment; (B) Pollen grains in 2017 experiment. Bars are fractions ± standard errors. Same letters above bars denote non-significant difference (P>0.05) between means. n.s. - not significant. Significance test was performed using GLM and contrasts analyses on relative number of pollen grains, normalized for each stigma by the mean number of pollen grains in all three stigmas of the same flower.

In *I. petrana*, 66 and 49 flowers were manipulated in 2016 and 2017, respectively. All stigmas received pollen grains in 2016, but in 2017, 51 (34%) did not receive any pollen grain. Number of pollen grains deposited on the stigmas was an order of magnitude larger than in *I. atropurpurea* in 2016 (Fig. 3 A), but in 2017 number of pollen grains was slightly less than in *I. atropurpurea* (Fig. 3 B).No significant treatment effect was found in 2016 (*F_2,195_*=0.60, *P*=0.242; Fig. 3 A). In 2017, however, we found significant higher number of pollen grains deposited on stigmas in flower units that manipulated in the low florivory treatment, compared to high florivory and control treatments (Contract analysis: P<0.001; Fig. 3 B).

In *Iris lortetii*, 11 flowers were manipulated in 2016. As in *I. petrana* in 2016, all stigmas received pollen and stigmas in units treated by high artificial florivory received pollen grains in a similar level as the control, untreated units, both higher than medium artificial florivory treatment (Fig. 3 A). However, this difference was not significant as well *(F_2,30_*=0.914, *P*=0.412).

### Fruit and seed sets

In *Iris atropurpurea*, 80 flowers, out of 96 flowers treated, were included in the final analyses in 2016, because 16 of the flowers were either not found or damaged. Of these, only 13 flowers set fruits, indicating extreme pollinator limitation and lack of pollinator visitations. Treatment did not affect either fruit-set or number of seeds (F*_2,75_*=0.69, *P*=0.501, and *F_2,8_*=0.26, *P*=0.774, respectively; Fig. 4 A and C). In both analyses flower size did not affect significantly (*P*=0.266 and *P*=0.554 for fruits and seeds, respectively). In 2017, 124 flowers were included in the experiment, of which four were damaged. Of the remaining 120 flowers, 23 flowers set fruits. As in 2016, no effect of the treatment was found, neither on fruit-set, nor on number of seeds (*F_2,116_*=0.37, *P*=0.695, and *F_2,19_*=0.95, *P*=0.404, respectively; Fig. 4 B and D). Similar to 2016, flower size as covariate did not affect fruit-set or seed-set (*P*=0.188 and *P*=0.857, respectively).

**Figure 4.**
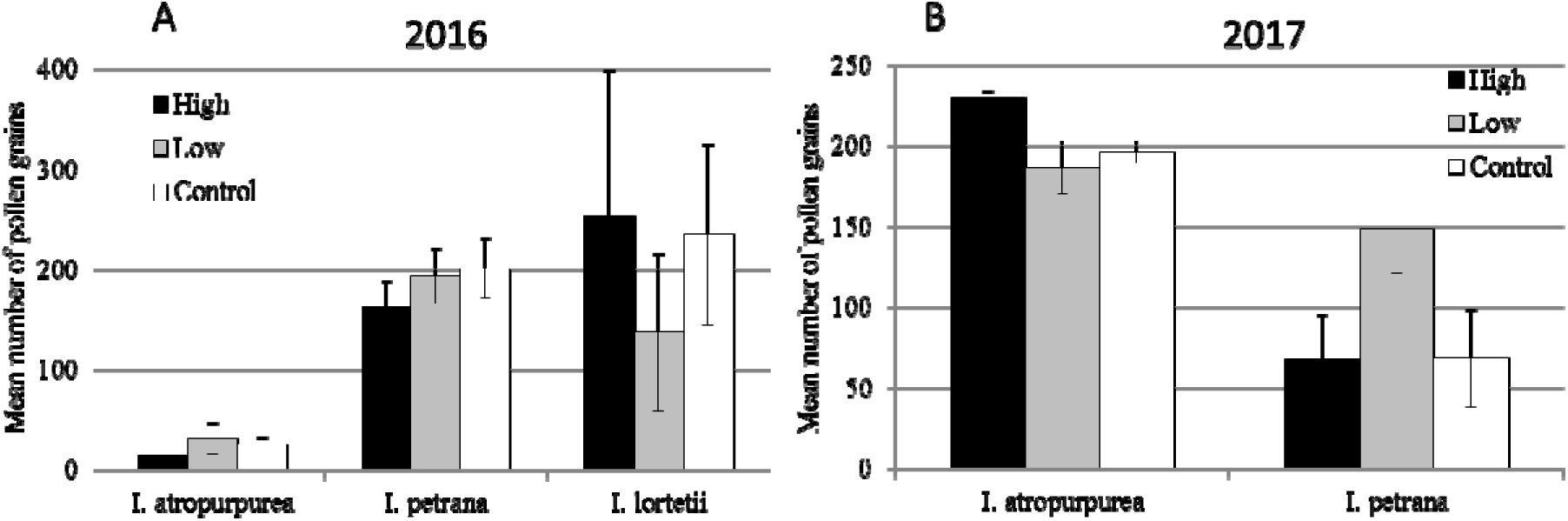
Fitness as a function of artificial florivory manipulations. (A) Fruit-set (fraction of flowers that set fruits) in three species in 2016 and (B) in two species in 2017. (C) Seed-set (mean number of seeds in a fruit) in 2016 and (D) in 2017. Bars are means ± standard errors.

In *I. petrana* population in Yeruham, 132 flowers were treated in 2016, but 29 flowers of all treatments were eaten by goats that entered the reserve illegally and ate wilting flowers and young fruits in the pre-dispersal stage. Of the remaining 103 treated flowers, 31 flowers set fruits. Flower length (before treatment) significantly affected fruit set (*F_1,98_*=5.09, *P*=0.026), but with no significant interaction with treatment (*F_2,96_*=0.26, *P*=0.770). Although control flowers produced almost twice fraction of fruits compared to florivory treatment (34% vs. 18%; contrasts analysis: P=0.022; Figure 4 A), this difference was not significant when controlled for flower size (*F*_2,98_=1.89, *P*=0.156). Number of seeds was not affected by treatment (*F_2,98_*=1.89, *P*=0.156, n=31; Fig. 4 C). Flower size did not affect seed-set (*F_1,26_*=1.23, *P*=0.278). In 2017, only six flowers were eaten or not found, out of 188 flowers treated. As in 2016, no effect of the treatment was found, neither on fruit-set, nor on number of seeds (*F_2,133_*=1.64, *P*=0.199, and *F_2,29_*=0.72, *P*=0.495, respectively; Fig. 4 B, D). As opposed to 2016, flower size as covariate did not affect fruit-set but did affect seed-set (*P*=0.472 and *P*<0.001, respectively). No interaction was found between flower size and florivory treatment in their effect on seed-set (*P*=0.463).

In the two sites of *Iris lortetii*, 61 out of 62 flowers treated (12 in Avivim and 50 in Malkiya) were found at the end of the season and included in the analyses. No significant difference was found among treatments (*F_2,57_*=0.21, *P*=0.811; Fig. 4 A). Flower size affected fruit-set (*F_1,57_*=4.42, *P*=0.40). Number of seeds was not significantly affected by treatment *(F_2,12_*=0.37, *P*=0.698; Fig. 4 C), and neither by flower size *(F_1,12_*=0.92, *P*=0.356).

## DISCUSSION

Florivory, the damage herbivores cause to floral organs, can affect fitness either directly by consuming pollen or ovules or by physiological costs, or indirectly, by reducing plant attraction signal for the pollinators (Burgess, 1991; McCall and Irwin, 2006). Here we tested for both direct and indirect effects of florivory on fitness by executing artificial florivory and measuring both fitness and pollination success. Our results do not fully support the hypothesis that florivory affects pollination success in the Royal Irises either directly or indirectly. Instead, we show that artificial damage to reproductive tissues in three species of the royal irises had inconsistent effect on pollen deposition, but no effect on fitness (Fig. 3, 4).

Florivory has small effect on pollinators’ behavior, but no overall effect on fitness. Limited cost in reducing pollinator’s services, no effect on fruit/seed-set. Adapted against florivory?

While numerous studies are concerned with the effect of florivory on plant fitness, controlled, artificial florivory was rarely performed. Most studies examined flowers that were naturally attacked by florivores (e.g., Meindl et al., 2013; Ruane et al., 2014; Eliyahu et al., 2015) or used experimental florivore removal or prevention (e.g., Krupnick et al., 1999; Theis and Adler, 2012; Althoff et al., 2013). Only a few studies implemented methods similar to ours, using cutting flowers to simulate florivory. These studies revealed mixed results. For example, Sõber et al. (2010) experimentally showed a correlation between extent of florivory and pollinator visitations at both population and plant level. On the other hand, Tsuji et al. (2016) found no evidence for pollinators discrimination against experimentally damaged flowers. Interestingly, mixed results can be found within the same system: Carper et al. (2016) found differences between heterostylous morphs in pollinator responses to artificial damage, and found no effect on fitness. Our study adds to the puzzle by providing yet another piece of evidence that florivory itself does not deter pollinators, nor reduces fitness.

Negative effect of florivory on pollination may act in two avenues. One possible effect of florivory on pollinator behavior is the deterring of pollinators from eaten flowers. This is achieved by either avoidance of flowers where florivore is visually detected (Kirk et al., 1995) or by a change in volatile compounds (Kessler et al., 2013). Another possible effect of florivory on pollination is mediated by the effect of the overall advertisement size of the flower and reducing visual signaling for the pollinators (Sánchez-Lafuente, 2007). This may reduce number of visits and lead to pollinator limitation or pollen limitation, which in turn reduces fruit-set and seed-set, respectively (Sapir et al., 2015). This study was conducted in natural populations of which at least one iris species, *Iris atropurpurea* experience strong pollinator limitation. It is likely that the effect of pollen limitation in this population obscures the effect of florivory (McCall, 2010). Thus, we propose that selection mediated by pollinator/pollen limitation is stronger than selection pressure exerted by florivory. A previous study that tested for pollen limitation in two *Iris* species showed that pollinator limitation provides conditions for pollinator-mediated selection (Lavi and Sapir, 2015). Although for *Iris petrana* we found no evidence for pollen limitation, still no effect of florivory was found, which contradicts this hypothesis. Our mixed results suggest that while pollinators may be a selection agent on flower traits (as in Lavi and Sapir, 2015), florivores do not act as selection agents because pollen limitation balances their effect (cf. Jogesh et al., 2017), but this connection cannot be generalized beyond our specific system.

Pollinators are thought to be the major selection agent on floral traits through their positive effect on fitness; this, however, was challenged by observations on contrasting effects of abiotic conditions or antagonistic biotic interactions (Herrera, 1996, reviewed in Strauss and Whittall, 2006). Strauss and Whittall (2006) proposed two scenarios of such mutualistic-antagonistic effect, in which overall selection acts as either directional or stabilizing on floral traits. Based on our results, we propose a third scenario, where florivory affects fitness at the same level as the mean effect of pollinators and regardless floral trait. In this case, pollinator-mediated selection will govern trait evolution, but the presence of florivory reduces or diminishes effect size (Figure 5). We speculate that florivory may not affect the direction of selection but the intensity of it. Thus, we suggest that testing for the net selection mediated by pollinators should control experimentally for the effect of florivores. In a follow-up experiment, we intend to test for net pollinator-mediated selection on floral color in the Royal Irises. Floral pigment concentration is expected to deter florivores and attract pollinators (de Jager and Ellis, 2014; McCall et al., 2013), and given the results of the current study we assume that florivory may not necessarily reduce fitness; instead, it is expected that a weak directional pollinator-mediated selection on floral color will be detected after controlling for florivory.

**Figure 5.**
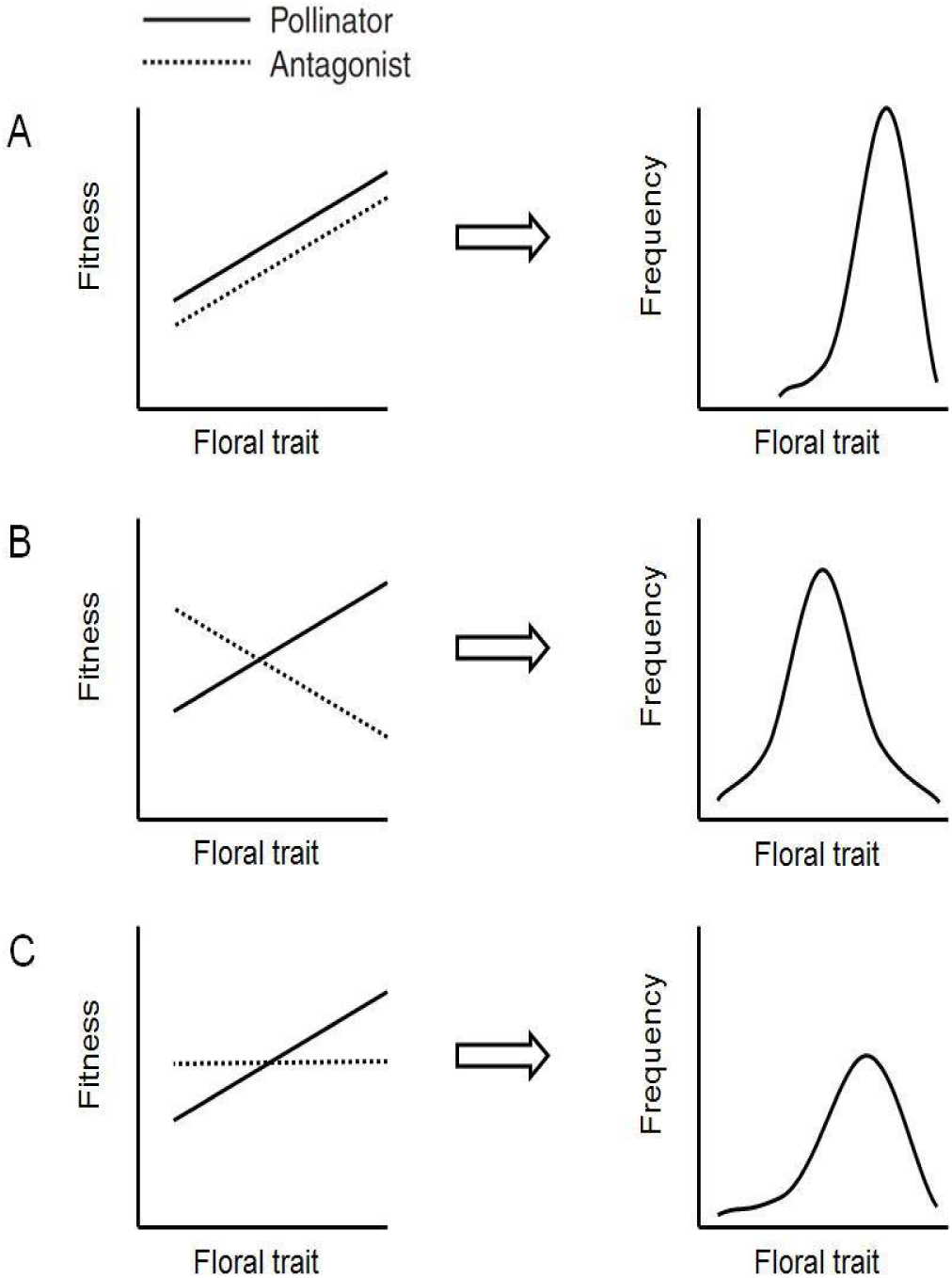
Hypothetical floral trait evolution as a function of selection by both mutualists (pollinators) and antagonists (florivores). (A) When pollinators and florivores exert selection in the same direction, the joint selection favors the same trait optimum. (B) When pollinators and florivores exert opposing selection on a trait, an intermediate trait optimum is favored. A and B are adapted from figure 7.1 in Strauss and Whittall (2006). (C) When florivores have no preference, or their effect is similar to mean fitness derived from pollinators’ effect, trait will have optimum fitness selected by pollinators like in A.

While few previous studies have detected only slight (or no) effect of florivory on pollination, it may be further hypothesized that the Royal Irises are unique and thus do not represent a general rule relevant to other species. The Royal Irises present a unique pollination syndrome in which pollination is performed by night sheltering male bees (Sapir et al., 2005; Watts et al., 2013). Previous studies suggest that floral display itself is not necessarily the major attractant for pollinators of the Royal Irises (Lavi and Sapir, 2015). While our florivory-like manipulations were performed on petals, it is likely that these manipulations did not affect pollinator choice because the shelter itself (the pollination tunnel) was not damaged. Because we have observed florivory of stigma and anthers (M. Ghara pers. obs.), we suspect that florivory may affect male and female fitness through consumption of reproductive organs, but we have yet to assess the cost of florivory on reproductive organs. Nonetheless, while natural florivory is widespread in all species and most populations of the Royal Irises in Israel, the estimated proportion of flowers of which pollination tunnels were eaten is rather small (M. Ghara, manuscript in preparation). Thus, our manipulation on petals accurately mimics natural florivory and our conclusions on the role of florivory in selection on flowers of the Royal Irises are valid.

Finally, our study provides an application to conservation. Of the three species studied, two *(Iris atropurpurea* and *I. lortetii)* are rare and endangered species (Sapir, 2016a, b). Understanding the relative contribution of biotic interactions to population dynamics may shed light on the factors affecting species survival in a way that may contribute for evidence-based management. The study presented here suggests that reduced mutualistic interactions, namely, pollination services, rather than antagonistic florivory, threatens the maintenance of positive population growth in the Royal Irises.

## CONCLUSIONS

Our experiments manipulations closely represent natural florivory in the Royal Irises. Florivores, largely considered as antagonists, do not seem to effect pollination in all the species we investigated. This is in contrast to some studies that found a negative effect of florivory on pollination where damage to flowers petals show reduction in fruit set. Perhaps, this is because of the unique pollination system in the Royal Irises where reward is shelter and the choice of shelter is carried out during the dusk and probably florivory cannot be truly assessed during those hours. Assessment of more study systems is thus required to arrive at a general conclusion of whether florivory affects pollination.

## ACKNOWLEDGEMENTS

The authors thank many students, volunteers and friends for assisting in field work in various capacities, especially Danya Ayehlet Cooper, Ilan Etam. And Yuval Shtrool. Prof. D. Eisikowitch for lab support and fruitful discussions as well as ideas. Dr. M Kate Gallagher provided critical comments on the manuscript. This study was partially supported by the Israeli Nature-Parks Authority and a grant from Israel Science Foundation (grant number 336/16) to YS. MG was supported by Israel Planning and Budgeting Commission (PBC) post-doctoral fellowship.

